# Phospholipid Scramblase-1 is required for efficient neurotransmission and synaptic vesicle retrieval at cerebellar synapses

**DOI:** 10.1101/2023.10.14.562324

**Authors:** Margherita Caputo, Daniela Ivanova, Sylvette Chasserot-Golaz, Frédéric Doussau, Anne-Marie Haeberlé, Sebahat Ozkan, Jason Ecard, Marie-France Bader, Nicolas Vitale, Michael A. Cousin, Petra Tóth, Stéphane Gasman, Stéphane Ory

## Abstract

Structural phospholipids are asymmetrically distributed at the plasma membrane, with phosphatidylethanolamine and phosphatidylserine (PS) virtually absent from the outer leaflet. This asymmetric lipid distribution is transiently altered during specific biological processes including calcium-regulated exocytosis. However, the impact of this transient remodeling of membrane asymmetry on presynaptic function remains unknown.

PhosphoLipid SCRamblase 1 (PLSCR1), a protein that randomizes phospholipid distribution between the two leaflets of the plasma membrane in response to calcium activation is an ideal candidate to alter this asymmetry. We therefore set out to determine the role of PLSCR1 in both neurotransmitter release and synaptic vesicle recycling by combining electron microscopy, optical live cell imaging of pHluorin probes and electrophysiology in cerebellar granule cells (GrC) from *Plscr1* knock-out mice (*Plscr1*^-/-^). We report that PLSCR1 is expressed in GrCs and that PLSCR1-dependent PS egress occurred at synapses in response to neuron stimulation. Furthermore, synaptic transmission is impaired at GrC *Plscr1*^-/-^ synapses and both PS egress and synaptic vesicle endocytosis are inhibited in *Plscr1*^-/-^ cultured neurons, demonstrating that PLSCR1 controls phospholipid asymmetry remodeling and synaptic vesicle retrieval following neurotransmitter release. Altogether, our data reveal a key role for PLSCR1 in synaptic vesicle recycling and provide the first evidence that phospholipid scrambling at the plasma membrane is a prerequisite for optimal presynaptic performance.

## Introduction

Structural phospholipids (PLs) at the plasma membrane are asymmetrically distributed by energy-dependent transporters, which move phosphatidylethanolamine (PE) and phosphatidylserine (PS) from the extracellular to the cytoplasmic leaflet (flippases) and, conversely, phosphatidylcholine (PC) from the cytoplasmic to the extracellular leaflet (floppases). PS and PE are therefore virtually absent from the outer leaflet of the plasma membrane (Clarke et al., 2020; Kobayashi and Menon, 2018; Zachowski, 1993). This homeostatic and asymmetric lipid distribution is disrupted in several biological processes resulting in PS exposure at the extracellular leaflet. Cell surface PS exposure is massive and irreversible during events such as apoptosis or platelet activation which is required for thrombosis (Bevers and Williamson, 2016). In contrast during regulated exocytosis in either immune or neuroendocrine cells, PS egress is transient and strictly restricted to sites of membrane fusion (Audo et al., 2017; Kato et al., 2002; Malacombe et al., 2006; Martin et al., 2000; Martin et al., 2000; Ory et al., 2013; Rysavy et al., 2014; Vitale et al., 2001).

Whether massive or limited, the loss of plasma membrane asymmetry is mainly due to the activation of phospholipid scramblases which randomize PLs across the plasma membrane (Daleke, 2003; Kobayashi and Menon, 2018). Phospholipid scramblase-1 (PLSCR1), originally isolated from erythrocytes, was the first scramblase identified able to reproduce lipid scrambling in response to Ca^2+^ increases when introduced into liposomes (Bassé et al., 1996; Zhou et al., 1997). PLSCR1 also controls PS egress in response to mast cell and chromaffin cell stimulation (Amir-Moazami et al., 2008; Kato et al., 2002; Ory et al., 2013; Smrž et al., 2008). However, preventing PS egress by suppressing PLSCR1 expression leads to different functional outcomes. For example, the release of secretory granule contents by exocytosis is reduced in mast cells (Amir-Moazami et al., 2008), but unaltered in chromaffin cells (Ory et al., 2013) which instead show defective compensatory endocytosis, preventing secretory granule protein retrieval (Ory et al., 2013). Therefore, the leaflet organization of the lipids at the plasma membrane and/or PLSCR1 control exocytosis and/or endocytosis in secretory cells. However, the mechanism of lipid scrambling by PLSCR1 is a currently a matter of debate. This is because PLSCR1 is a single pass transmembrane protein that cannot form a channel to transport PLs, and the deletion of *Plscr1* gene in mice does not alter plasma membrane PS egress either in activated platelets or in response to apoptosis (Zhou et al., 2002). Nonetheless, the identification of additional scramblases, namely TMEM16F and XKR-8 controlling extracellular PS exposure during platelet activation (Suzuki et al., 2010; Yang et al., 2012) and apoptosis (Suzuki et al., 2013), respectively, provides evidence that, despite their common ability to collapse lipid asymmetry, scramblases can regulate distinct mechanisms at the plasma membrane.

Neuronal communication relies on the release of neurotransmitters at synapses, which occurs by Ca^2+^-dependent exocytosis of synaptic vesicles (SVs), resulting in the insertion of SV membrane into the presynaptic plasma membrane. To support high rates of release during synaptic transmission, SV components must be retrieved rapidly by compensatory endocytosis to preserve plasma membrane homeostasis, to clear exocytic sites for the recruitment of release-ready SVs and to replenish the releasable SV pool (Chanaday et al., 2019; Maritzen and Haucke, 2018). Activity-dependent SV fusion and retrieval are spatially and temporally coupled within nerve terminals and dysfunction in either process is increasingly associated with pathologies including epilepsy, autism or intellectual disability (Bonnycastle et al., 2020). The molecular mechanisms coupling endocytosis to exocytosis have been extensively studied in neurons (Bolz et al., 2023; Maritzen and Haucke, 2018). However, although exocytosis and endocytosis imply successive reorganization of PLs belonging to separate membrane compartments, little attention has been paid to the role of plasma membrane PLs in these events.

The present study sought to determine whether SV recycling relied on PLSCR1, and whether neurotransmission was influenced by its loss. Through a combination of approaches including immunofluorescence on primary cerebellar cultures, electron microscopy and electrophysiology on acute cerebellar slices from both wild-type (*Plscr1*^+/+^) and *Plscr1* knock-out mice (*Plscr1*^-/-^), we revealed that PLSCR1-dependent PS egress at synapses is essential for maintaining efficient neurotransmission. Moreover, optical live cell imaging of pHluorin probes revealed that compensatory endocytosis was altered in PLSCR1^-/-^ neurons. Taken together, our data unveil a pivotal role of PLSCR1 in mediating stimulation-dependent PS egress at excitatory synapses. Furthermore, our study provides the first evidence of the impact of PLSCR1-dependent PS egress on facilitating SV recycling and sustaining synaptic transmission during periods of high frequency stimulation.

## Material and Methods

### Animals and genotyping

Neuronal cells and brain tissues were obtained from *Plscr1*^-/-^ and wild type mice. *Plscr1*^+/−^ mice were purchased from CDTA (Cryopréservation, Distribution, Typage et Archivage animal) housed and raised at Chronobiotron UMS 3415. All mice were bred, handled, and maintained in agreement with European council directive 86/609/EEC and resulting French regulation. Animal work in Edinburgh was performed in accordance with the UK Animal (Scientific Procedures) Act 1986, under Project and Personal License authority and was approved by the Animal Welfare and Ethical Review Body at the University of Edinburgh (Home Office project license to M. Cousin – 70/8878). Animals were killed by schedule 1 procedures in accordance with UK Home Office Guidelines. Animals used in France were handled according to French regulations. P6 to P7 mice pups were killed by decapitation and adult mice were euthanized by cervical dislocation. *Plscr1^+/+^* and *Plscr1*^-/-^ mice were maintained as heterozygous breeding pairs and genotyped by duplex PCR using 5’-CTACACTGACCTTTAATCAGAGCAG-3’, 5’-CCATGTCTGCCCAAGTTCACTCTC-3’, and 5’-GCAGCGCATCGCCTTCTATC-3’ primers to detect the presence of a 261 bp or a 311 bp fragment for the wild-type or mutant allele respectively. The PCR program was set at 95°C for 3 min; 30 cycles (95°C for 30 sec, 60°C for 30 s,72°C for 30 sec) followed by 10 min elongation at 72°C.

### Cell culture, transfection and plasmids

Cerebellar granule cultures were prepared as previously described (Cheung and Cousin, 2011). Briefly, primary cultures of cerebellar neurons were prepared from the cerebella of 6-7 days old C57BL/6J *Plscr1*^-/-^ pups and their littermate controls. After removal, cerebella were placed in HH solution (Table 1), minced up with tweezers and then incubated into the Trypsin solution (2T; Table 1) at 37°C for 20 min. After digestion with trypsin, an equal volume of Neutralization solution (N; Table 1) was added to the suspension and the sample was centrifuged at 150 × *g* for 60 s. The supernatant was removed and the pellet suspended in 1 mL of N solution before being triturated using 1000 µL pipette until the solution reached homogeneity. The cell suspension was centrifuged at 340 × *g* for 2 min and the pellet resuspended in 1.5 mL of DMEM/FCS medium (DFM)(Table 1) and plated as one spot/well at a density of 5–10 × 10^6^ cells/coverslip coated with poly-D-lysine (20 μg/ml) diluted in boric acid (100 mM, pH 8.5). After 1 hour, wells were flooded with Neurobasal growth medium (NKB) (Table 1) containing 25 mM KCl. The following day, cultures were further supplemented with 1 μm cytosine β-D-arabinofuranoside to inhibit glial cell proliferation. 4 or 5 days after seeding, cells were transfected with Lipofectamine 2000 as described by Nicholson-Fish et al. (2015). Briefly, cells were preincubated in 2 ml of MEM (Thermo Fisher) in 10% CO_2_ for 30 min at 37°C, and then transfected for 2 hours with a complex containing 2 μL of Lipofectamine and 1 μg of the indicated plasmids/well. Cells were subsequently washed with MEM before replacement with conditioned Neurobasal media. Cells were imaged 48 h post-transfection.

**Table 1:** Solutions for CGN preparation.

|  |  |
| --- | --- |
| HH Solution | HBSS $\text{Ca}^{2+}/\text{Mg}^{2+}$ free (Sigma, H6648) supplemented with 5 mL of 1M sterile Hepes pH 7.3 |
| Trypsin Solution (2T) | 1 mL of sterile 5mg/mL trypsin in 9 mL of HH to make final 0.05% solution |
| DFM medium | DMEM high glucose (Thermo Fisher, 41966029) supplemented with 10%FCS and 1% Pen/Strep |
| Neutralization Solution (N) | 1 mL of sterile DNase containing 1200 U in 19 mL of DFM to make final 60U/mL solution |
| NBK medium | Neurobasal (Thermo Fisher 21103049) supplemented with 2% B27, 0.25% 200 mM Glutamine and 1% Pen/Strep |

Synaptophysin-pHluorin gifted from Pr Lagnado (University of Sussex, Brighton, UK) and G FP-PLSCR1 plasmids were previously described (Granseth et al., 2006; Ory et al., 2013). The mCherry-PLSCR1 constructs were obtained by PCR using the mouse *Plscr1* as a template. Amplified fragments were subcloned in pmCherry-C1 vectors between BglII and EcoRI restricti on sites using forward 5’-CAGATCTGAAAACCACAGCAAGCAAAC-3’ and reverse primer 5’ –GG AATTCTTACTGCCATGCTCCTGATC-3’.

### Immunofluorescence, Confocal microscopy and Image analysis

Neurons were fixed with ice cold 4% (w/v) paraformaldehyde in phosphate buffered saline (PBS) for 10 min at room temperature and permeabilized with 0.1% v/v Triton x-100 in PBS for 4 min. The cells were washed with PBS, blocked with 3% BSA in PBS and 5% Goat Serum (Sigma, #G9023) for 1h before being incubated with primary antibodies in 3% BSA in PBS for 1h. Then cells were labelled with secondary antibodies coupled to fluorescent Alexa Fluor dyes (see Table 2 for antibodies list and references). Actin was stained with Tetramethyl-rhodamineB coupled Phalloidin (TMR-phalloidin, Table2) and nuclei were stained with 1 µg/mL Hoechst (Thermo Scientific, 33342). Cells were observed under a confocal microscope equipped with continuous laser emitting at 405, 488, 561 and 633 nm (SP5, Leica Microsystems), using a 63× objective (NA 1.4).

**Table 2.**

| Reagents | Company | Reference | Dilution | RRID |
| --- | --- | --- | --- | --- |
| Rabbit anti Synaptotagmin1 | Synaptic systems | 105103 | 1/1000 | AB_11042157 |
| Guinea pig anti vGlut1 | Synaptic systems | 135304 | 1/500 | AB_887878 |
| Mouse anti Synaptophysin | Sigma | S5768 | 1/500 | AB_477523 |
| Rabbit anti-GFP | Abcam | ab290 | 1/1000 | AB_303395 |
| Mouse anti actin | Sigma | A5441 | 1/1000 | AB_476744 |
| Mouse anti PLSCR1 | Proteintech | 11582-1AP | 1/500 | AB_2165659 |
| Mouse anti PSD95 | Thermofisher scientific | MA1-045 | 1/500 | AB_325399 |
| HRP-conjugated goat anti-mouse and anti-rabbit | PerBio | 31430, 31460 | 1/50000 | AB_228307, AB_228341 |
| Alexafluor647®-conjugated goat anti guinea pig, Alexafluor555®-conjugated goat anti rabbit, | Thermofischer scientific | A21450, A21428 | 1/1000 | AB_2633280, AB_26332276, AB_2534120 |
| Gold particle-conjugated anti-rabbit and anti guinea pig | Aurion | 810 144, 825.011 | 1/100 | N/A |
| TMR-Phalloidin | Sigma | P1951 | 1/1500 | AB_2315148 |
| AnxA5-15 nm gold conjugated | Biobyte | Orb170087 | 1/50 | N/A |
| AnxA5-AlexaFluor 647 | Biolegend | 640911 | 1/50 | N/A |
| Hoechst | Thermoscientific | 33342 | 1/10000 | N/A |
| Picrotoxin | AbCam | Ab120315 | 100 µm in ACSF | N/A |
| CGP52432 | AbCam | Ab120330 | 10 µm in ACSF | N/A |
| D-AP5 | AbCam | Ab120003 | 100 $\mu\text{m}$<br>in ACSF | N/A |
| AM251 | AbCam | Ab120088 | 1 $\mu\text{m}$ in<br>ACSF | N/A |
| JNJ162259685 | Tocris | 2333 | 2 $\mu\text{m}$ in<br>ACSF | N/A |

Annexin-A5 (AnxA5) staining was performed as described before (Ory et al., 2013). Cerebellar granule cells were hyperpolarized for 10 min in Imaging Buffer (170 mM NaCl, 3.5 mM KCl, 0.4 mM KH_2_PO_4_, 20 mM TES (N-tris (hydroxyl-methyl)-methyl-2-aminoethane-sulfonic acid), 5 mM NaHCO_3_, 5 mM glucose, 1.2 mM MgCl_2_, and 1.3 mM CaCl_2_, pH 7.4). Neurons were then maintained for 10 min at 37°C in the presence of AlexaFluor-647-conjugated AnxA5 (1/50, Biolegend) in resting solution (Locke’s solution:140 mM NaCl, 4.7 mM KCl, 2.5 mM CaCl_2_, 1.2 mM KH_2_PO_4_, 1.2 mM MgSO_4_, 11 mM glucose, and 15 mM HEPES, pH 7.2; resting) or in stimulation solution containing 59 mM K^+^ (Locke’s solution containing 59 mM KCl and 85 mM NaCl). Neurons were then fixed and counterstained for 30 min with TMR-phalloidin. To identify PS egress sites, cells were incubated in culture medium containing 30 mM KCl and both AlexaFluor-647-conjugated AnxA5 and anti-Synaptotagmin1 (Syt1) antibody directed against the luminal domain. Cells were then fixed and Syt1 antibodies revealed with a goat anti-rabbit antibody conjugated to Alexa Fluor 555. AnxA5 and Syt1 staining were observed under confocal microscope (SP5, Leica Microsystems). Image analyses were performed with Icy software. HKmeans thresholding method (Manich et al., 2020) was used to segment actin staining and mean AnxA5 intensity was computed in the resulting region of interest (ROI) delineating the neuronal shape. AnxA5 mean intensity was obtained from 10 fields of view per each independent experiments (n=3).

### Preparation of Synaptosomal Fraction

Synaptosomal fractions were prepared as previously described (Dunkley et al., 2008a). Briefly, cerebella from adult mice were collected in an isotonic Sucrose/EDTA buffer (0.32 M Sucrose, 1 mM EDTA, 5 mM Tris, pH 7; Homogenizing Buffer) and dissociated by applying 13 even strokes with a dounce homogenizer. After centrifugation of the homogenates (1000 *g*, 10 min, 4°C), the supernatant (S1, Crude extract) was kept on ice and the pellet was resuspended in homogenizing buffer, subjected to additional 17 even strokes and centrifuged (1000 *g*, 10 min, 4°C). The pellet was discarded and the supernatant S2 was pooled with S1. Protein concentration was measured and adjusted with ice-cold homogenizing buffer to 4-5 mg/mL. Crude extract was loaded onto a discontinuous Percoll gradient (3%, 10%, 15% and 23% Percoll (vol/vol)) and centrifuged at 31,000g for 5 min, at 4°C. Each fraction was individually recovered and fractions 3 and 4 enriched in synaptosomes were pooled. Synaptosomes were then diluted in ice-cold homogenizing buffer and centrifuged at 20,000g for 30 min at 4°C to concentrate the synaptosomes into the pellet and remove Percoll. Synaptosomes were resuspended in lysis buffer (Cell Extraction Buffer: 10 mM Tris, pH 7.4, 100 mM NaCl, 1 mM EDTA, 1 mM EGTA, 1 mM NaF, 20 mM Na_4_P_2_O_7,_ 2 mM Na_3_VO_4_, 1 % Triton X-100, 10 % glycerol, 0.1 % SDS, 0.5 % deoxycholate; Invitrogen, #FNN0011) supplemented with protease and phosphatase inhibitor cocktail (Sigma-Aldrich, # P8340) and further subjected to Western Blot analysis.

### Western Blotting

Cells and tissues were lysed in cell extraction buffer with protease and phosphatase inhibitor cocktail (P8340, Sigma-Aldrich). Cell lysates were cleared at 20,000g for 10 min at 4°C and protein concentration determined using BioRad protein assay. Proteins (20 μg total) were separated on Novex 4-12% Bis-Tris gels (ThermoFisher Scientific) and transferred to nitrocellulose membrane (BioRad, #1704156). Blots were blocked for 1 h at room temperature in Tris-buffered saline containing 5% (w/v) milk powder (fat free) and 0.1% Tween-20 (TBST; 0.1% Tween, 150 mM NaCl, 10 mM Tris-HCl, pH 7.5) and probed with the anti-PLSCR1, anti-Synaptotagmin1 or anti-PSD95 antibodies (see Table 2). After three washes, blots were incubated with the corresponding secondary antibodies coupled to HRP (Table 2). Detection was carried out with Prime Western Blotting System (ThermoFisher Scientific) and immunoreactive bands were imaged using Amersham Imager 680 RGB camera System (GE healthcare Life Sciences). Values were normalized to the corresponding β-actin protein levels.

### Acute slice preparation and electrophysiological recordings

Slice preparation and electrophysiology recording were performed as previously described (Doussau et al., 2017). Acute horizontal cerebellar slices were prepared from male and female C57Bl/6 mice of both genotype (*Plscr1*^+/+^ and *Plscr1*^-/-^) aged 18-30 days. Mice were anesthetized by isoflurane inhalation (4%) and decapitated. The cerebellum was dissected out in ice-cold artificial cerebrospinal fluid (ACSF) bubbled with carbogen (95% O_2_, 5% CO_2_) and containing 120 mM NaCl, 3 mM KCl, 26 mM NaHCO_3_, 1.25 mM NaH_2_PO_4_, 2.5 mM CaCl_2_, 2 mM MgCl_2_, 10 mM glucose and 0.05 mM minocyclin. Slices were then prepared (Microm HM650V) in an ice-cold solution containing 93 mM *N*-Methyl-D-Glucamine, 2.5 mM KCl, 0.5 mM CaCl_2_, 10 mM MgSO_4_, 1.2 mM NaH_2_PO_4_, 30 mM NaHCO_3_, 20 mM HEPES, 3 mM Na-Pyruvate, 2mM Thiourea, 5 mM Na-ascorbate, 25 mM D-Glucose and 1 mM Kynurenic acid (Zhao et al., 2011). Slices 300 µm thick were maintained in bubbled ASCF medium (see above) at 34°C until their use for experiments.

After at least 1 h of recovery at 34°C, a cerebellar slice was transferred to a recording chamber. In order to block inhibitory transmission, postsynaptic plasticities, GABA_B_ and endocannabinoid signaling, slices were continuously perfused with bubbled ACSF containing blockers of GABA_A_, GABA_B_, NMDA, CB1 and mGluR1 receptors. To do so, the following antagonists were added (see Table2 for references): 100 µM picrotoxin, 10 µM CGP52432 (3-[[(3,4-Dichlorophenyl)-methyl]amino]propyl(diethoxymethyl)phosphinic acid), 100 µM D-AP5 (D-(-)-2-Amino-5-phosphonopentanoic acid) and 1 µM AM251 (1-(2,4-Dichlorophenyl)-5-(4-iodophenyl)-4-methyl-N-(piperidin-1-yl)-1H-pyrazole-3-carboxamide) and 2 µM JNJ16259685((3,4-Dihydro-2H-pyrano[2,3-b]quinolin-7-yl)-(cis-4-methoxycyclohexyl)-methanone). Recordings were made at 34°C in Purkinje cells (PCs) located in the vermis. PCs were visualized using infrared contrast optics on an Olympus BX51WI upright microscope. Whole-cell patch-clamp recordings were obtained using a Multiclamp 700A amplifier (Molecular Devices). Pipette (2.5-3 MΩ resistance) capacitance was cancelled and series resistance (R_s_) between 5 and 8 mΩ was compensated at 80%. R_s_ was monitored regularly during the experiment and the recording was stopped when R_s_ changed significantly (> 20%). PCs were held at −60 mV. The intracellular solution for voltage-clamp recording contained 140 mM CsCH_3_SO_3_, 10 mM phosphocreatine, 10 mM HEPES, 5 mM QX314-Cl, 10 mM BAPTA, 4 mM Na-ATP and 0.3 mM Na-GTP. Beams of parallel fibers were stimulated extracellularly using a monopolar glass electrode filled with ACSF, positioned at least 100 µm away from the PC to ensure a clear separation between the stimulus artifact and EPSCs. Pulse trains were generated using an Isostim A320 isolated constant current stimulator (World Precision Instruments) controlled by WinWCP freeware (John Dempster, Strathclyde Institute of Pharmacy and Biomedical Sciences, University of Strathclyde, UK). The synaptic currents evoked in PCs were low-pass filtered at 2 KHz and sampled at 20 to 50 KHz (National Instruments).

### Cerebellum slices and plasma membrane sheets preparation for transmission electron microscopy

Wild-type (*n* = 3) and *Plscr1*^-/-^ (*n*=3) mice were anesthetized with a mixture of ketamine (100 mg/kg) and xylazine (5 mg/kg) and transcardiacally perfused with 0.1 M phosphate buffer, pH 7.3, containing 2% paraformaldehyde and 2.5% glutaraldehyde. The 2-mm-thick slices were cut from the cerebellum and postfixed in 1% glutaraldehyde in phosphate buffer overnight at 4°C. The slices were then immersed for 1 h in OsO4 0.5% in phosphate buffer. The 1 mm^3^ blocks were cut in the cerebellum, dehydrated, and processed classically for embedding in araldite and ultramicrotomy. Ultrathin sections were counterstained with uranyl acetate. Fields of view were randomly selected in ultrathin sections from several blocks (1 section/block) from each mouse.

Membrane sheets were prepared and processed as described previously (Delavoie et al., 2021; Gabel et al., 2015). In brief, carbon-coated Formvar films on nickel electron grids were inverted onto unstimulated or stimulated GrC (Locke buffer containing 50 mM KCl) incubated with gold-conjugated AnxA5 for 10 min. To prepare membrane sheets, pressure was applied to the grids for 25 s, then grids were lifted so that the fragments of the upper cell surface adhered to the grid. These membrane sheets were then fixed in 2% paraformaldehyde for 10 min at 4°C, blocked in PBS with 1% BSA and 1% acetylated BSA and incubated with antibodies anti-GFP and anti-v-Glut1 (see Table 2) overnight at 4°C. Then the membranes were washed 6 times with PBS and incubated 3 h with 25 nm gold particle-conjugated goat anti-rabbit IgG and 10nm gold particle-conjugated goat anti-guinea pig IgG (Table 2). These membrane portions were fixed in 2% glutaraldehyde in PBS, postfixed with 0.5% OsO4, dehydrated in a graded ethanol series, treated with hexamethyldisilazane (Sigma-Aldrich) and air dried. Transmission electron microscope (7500; Hitachi) equipped with camera (C4742-51-12NR; Hamamatsu Photonics) were used for acquisition.

### Morphometric analysis of electron microscopy slices

SVs, pre and post-synaptic membranes and boutons were manually selected or delineated using Icy bioimaging software. The density of SVs in the boutons and the shortest distance between SV and the pre-synaptic membrane were calculated. Synapse density per field were also calculated and the mean distance of synaptic cleft were computed.

### SynaptopHluorin live cell imaging

Imaging was performed with DIV6 to DIV7 cerebellar granule neurons as reported by (Nicholson-Fish et al., 2015). 48 to 72 hours post transfection, GrC were removed from culture medium and repolarized in Imaging Buffer (see above) for 15 min. Coverslips were then mounted in an imaging chamber (Warner RC-21BRFS) supplied with a pair of embedded platinum wires allowing connection to a low impedance field stimulator (Digitimer, D330 MultiStim System). The chamber was placed on the stage of an inverted microscope (Zeiss AxioObserver Z1) equipped with either a sCMOS camera (Orca-FLASH4, Hamamatsu) or a Zeiss AxioCam506 and a control of focus (Zeiss Definite Focus) to prevent drift. During recording, cells were continuously perfused with Imaging Buffer at 37°C. Synaptophysin-pHluorin (Syp-pH) transfected neurons were visualized with a Zeiss Plan Apochromat x40 oil-immersion objective (NA 1.4) at 475 +/- 28 nm excitation wavelength using 525 +/- 50 nm emission filter (LED light source SpectraX, Lumencor) or exciter 450-490 nm, beam splitter 495 nm, emitter 500-550 nm, Colibri 7 LED light source (Zeiss)). When specified, cotransfected mCherry expression was visualized using 555 +/- 28 nm excitation and 605 +/- 70 nm emission filters or exciter 538-652 nm, beam splitter 570 nm, emitter 570-640 nm. Neurons were stimulated with a train of action potentials (400 action potentials delivered at 40 Hz; 100 mA, 1-ms pulse width) and the acquisition sequence driven by MetaMorph Software (Molecular Devices) or Zeiss Zen2 software at 5 Hz frequency. At the end of the recording, cells were challenged with alkaline imaging buffer (50 mM NH_4_Cl substituted for 50 mM NaCl) to reveal total pHluorin fluorescence. Quantification of the time-lapse series was performed using the Time Series Analyzer plugin for ImageJ and only synapses that responded to action potential stimulation were selected for the analysis. A circular region of interest (ROI, 1.8 μm diameter) was drawn around each spot characterized by a sudden rise in fluorescence. ROI was centered on the maximum fluorescence of the spot. The pHluorin fluorescence change in each spot was calculated as FΔ/F_0_, and n refers to the number of individual cells examined. Statistical analyses were performed using Microsoft Excel and GraphPad Prism software.

#### Statistical analysis

Analysis was performed by using GraphPad Prism (2018). All data passed normality test (Shapiro–Wilk test) and variance equality. A Student’s t-test was performed for comparison between two datasets. Data were analysed by one-way analysis of variance (Anova) with Sidak’s multiple comparison test when greater than two datasets. To compare fluorescence response over time or data with more than one variable, two-way Anova with Tukey’s multiple comparison *post hoc* test was performed. All data are reported as mean ± standard error of the mean (SEM). To attest for similar SVs distribution, the Kolmogorov-Smirnov test was applied. Significance was set as *P < 0.05, **P < 0.01, ***P < 0.001. ns, non-significant

#### Databases

Preliminary information about *Plscr1* distribution in the mouse brain were obtained from the Allen brain atlas (https://mouse.brain-map.org/experiment/show/632487) and the Human brain atlas databases (https://www.proteinatlas.org/ENSG00000188313-PLSCR1/brain)

## Results

### PLSCR1 is expressed in cerebellar granular cells and localizes at synapses

PLSCR1 is expressed in a wide range of tissue including the brain (Wiedmer et al., 2000; Zhou et al., 2005). Mining in mouse brain databases revealed that the *Plscr1* transcript is barely detectable by *in situ* hybridization (Allen brain atlas, Lein et al., 2007). However, it is expressed in the olfactory bulb, the cerebellum, the pons and medulla when using the more sensitive next generation sequencing (Human Protein Atlas, Sjöstedt et al., 2020). To validate the expression profile of the PLSCR1 gene product in a specific area of the mouse brain, we dissected different brain areas to perform Western blots on tissue extracts from *Plscr1*^+/+^ adult mice. As negative control, we used cerebellar extracts from *Plscr1*^-/-^. We confirmed that PLSCR1 is abundant in the cerebellum, with expression also observed in the olfactory bulb and midbrain albeit at lower levels. However, expression is barely detectable in all the other brain regions investigated (Figure 1A). We therefore focused our studies on the cerebellum, which displayed the highest expression of PLSCR1 in mouse brain.

**Figure 1:**
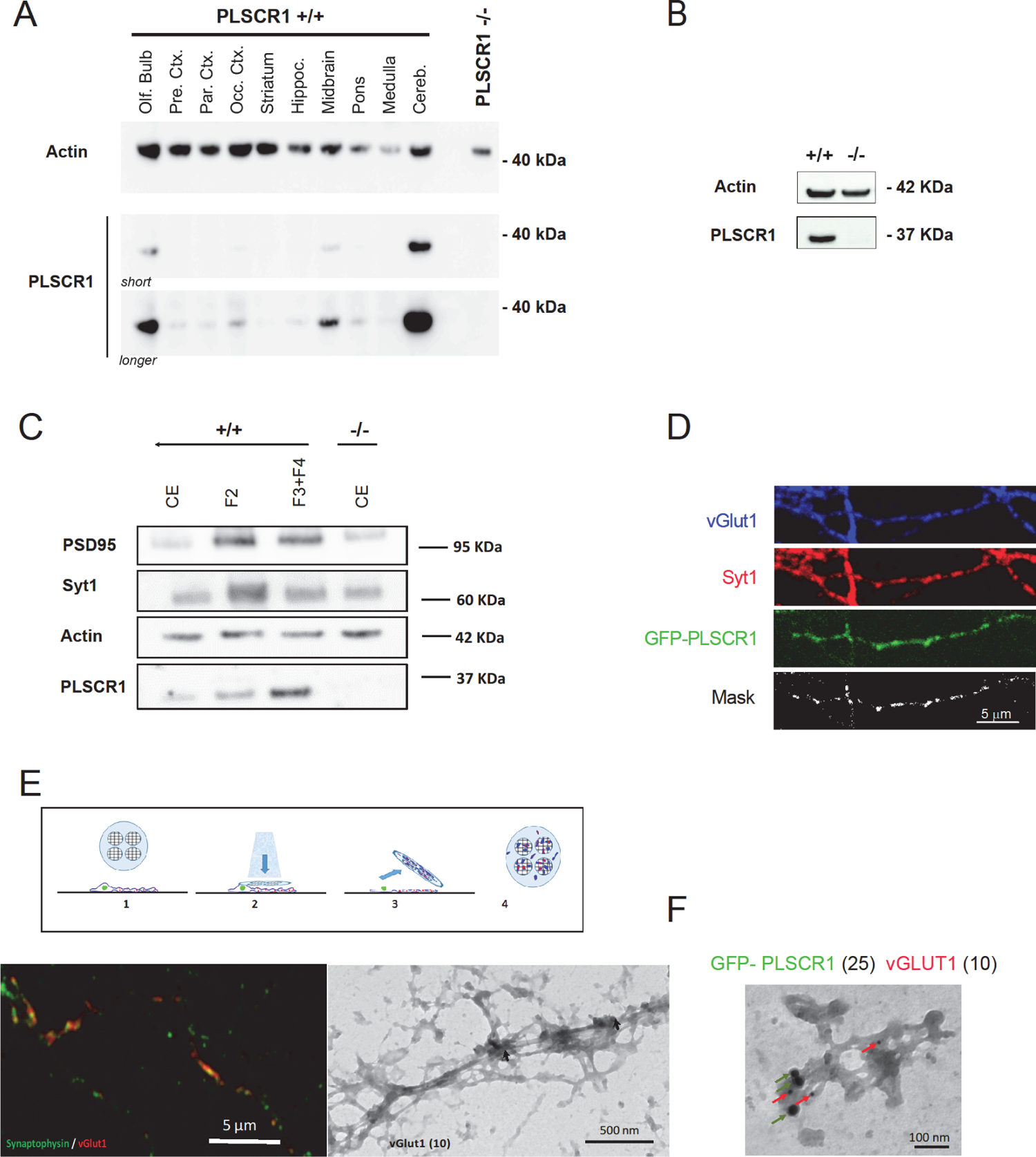
PLSCR1 is enriched at synapses of GrC neurons of the cerebellum. **A**, immunodetection of PLSCR1 protein by western blot in the olfactory bulb (Olf. Bulb), cortex (prefrontal (Pre. Ctx.), parietal (Par. Ctx.) and occipital (Occ. Ctx.), striatum, hippocampus (Hippoc.), midbrain, pons, medulla and cerebellum (Cereb.) from PLSCR1^+/+^ mice. As a control, cerebellum of PLSCR1^-/-^ mice was used (PLSCR1^-/-^). Blot with short and longer exposure (long) are shown. Actin is shown as a loading control. **B**, immunodetection of PLSCR1 protein by western blot in cultured GrC neurons. **C**, PLSCR1 is enriched in synaptosomes. Cerebella were homogenized, cell body removed by centrifugation and the resultant supernatant layered on a discontinuous Percoll gradient. Fractions were collected and probed for PLSCR1, the presynaptic marker Syt1 and the postsynaptic marker PSD95. F2 corresponds to the fraction enriched in membranes and fractions F3 and F4 enriched in synaptosomes were pooled and compared to crude extract (CE) loaded on gradient. **D**, confocal microscopy of GrC neurons transfected with expression vector coding for GFP-PLSCR1 and labeled for vGlut1 and Syt1. Mask of synapses containing the three markers is shown. **E**, principle of plasma membrane sheet preparation. *Top:* carbon-coated Formwar films on nickel EM grid (1) were inverted onto neurons grown on poly-L-lysine-coated coverslips and pressure is applied to the grid (2). Grids are then lifted (3), leaving fragments of plasma membrane (blue) and synapses containing docked SVs (red) on the grids (4)*. Bottom left:* confocal images of GrC plasma membrane sheets stained for vGlut1 (red) and synaptophysin (green). *Bottom right:* representative electron micrograph of GrC plasma membrane sheets labeled for vGlut1 (10nm gold particles, arrow). **F**, electron micrograph of plasma membrane sheets prepared from *Plscr1*^-/-^ GrC neurons expressing GFP-PLSCR1. Immunolabeling of GFP and vGlut1 were revealed with 25 nm (green arrows) and 10 nm (red arrows) gold particles respectively.

Since GrC account for 95% of cerebellum cells (D’Mello et al., 1993; Krämer and Minichiello, 2010), we first probed PLSCR1 expression in primary cultures of GrCs displaying high homogeneity. Western blot experiments revealed that PLSCR1 was expressed in GrCs cultured from *Plscr1^+/+^*mice but, as expected, absent from *Plscr1*^-/-^ GrCs culture (Figure 1B). We next analyzed PLSCR1 subcellular distribution. First, we prepared synaptosomes from mouse cerebellum to determine whether PLSCR1 was localized to synaptic terminals (Dunkley et al., 2008b). Compared to crude brain homogenate, PLSCR1 is enriched in synaptosomal fractions containing the post- and pre-synaptic markers PSD95 and Synaptotagmin 1 (Syt1), respectively (Figure 1C). To further investigate the presence of PLSCR1 at synapses, we performed immunofluorescence on GrC. Because of the lack of reliable antibodies to detect endogenous PLSCR1 by immunofluorescence, we exogenously expressed GFP-tagged PLSCR1 and compared its distribution with specific markers of the pre-synaptic terminal, the vesicular glutamate transporter1 (vGlut1) and Syt1. GFP-PLSCR1 was found in GrC axons and colocalized with vGlut1 and Syt1, indicating that it is also present at the synapse (Figure 1D). To further explore PLSCR1 localization, we performed immunogold electron microscopy (EM) analysis of native plasma membrane sheets of GrCs (Figure 1E). Sheets were obtained by tearing off the plasma membrane at the dorsal surfaces of cells to examine, at high resolution, the associated molecules (Gabel et al., 2015; Wilson et al., 2000).

Synaptic components were retained by this method, since confocal analysis of recovered membrane pieces were stained for Synaptophysin and vGlut1. Electron microscopy analysis using secondary antibodies coupled to 10 nm gold particles, showed vGlut1 clusters on membrane patches of various size (Figure 1E). Plasma membrane sheets from *Plscr1*^-/-^ neurons expressing GFP-PLSCR1 were analyzed. Secondary antibodies coupled to 10- and 25-nm gold particles were used to reveal both the expression of GFP-PLSCR1 and vGLUT1 respectively (Figure 1F). Gold particles of different size were found distributed over most membrane patches. In addition, GFP-PLSCR1 was localized close to vGlut1, with the majority of membrane patches containing vGlut1 also containing GFP-PLSCR1 (73% ± 13%; n=5 experiments). Altogether, our data indicate that GFP-PLSCR1 is associated to the plasma membrane of GrCs and is most likely enriched at the pre-synaptic terminal were vGlut1-positive SVs are located.

To determine whether *Plscr1* deletion could modify synaptic organization and/or characteristics, we performed morphometric analysis of sections from *Plscr1*^-/-^ and *Plscr1*^+/+^ cerebellum observed by transmission electron microscopy (Figure 2A). GrC to Purkinje cell (PC) synapses from *Plscr1*^-/-^ mice showed no significant differences in either synapse density per slice or SV density in synaptic boutons compared to *Plscr1*^+/+^ mice (Figure 2B). We measured the minimal distance separating each SV from the synaptic membrane facing the post-synaptic density, but found no significant differences in either the distribution of SVs (Figure 2C) or the number of docked SVs (Figure 2D). We did however note a slight decrease in synaptic length (Figure 2E). The synaptic cleft, measured as the space separating the pre- and post-synaptic membranes, remained unchanged (Figure 2E). Consequently, the absence of PLSCR1 has negligible impact on the structural organisation of GrC-PC synapses, suggesting that PLSCR1 does not influence developmental processes in the cerebellum.

**Figure 2:**
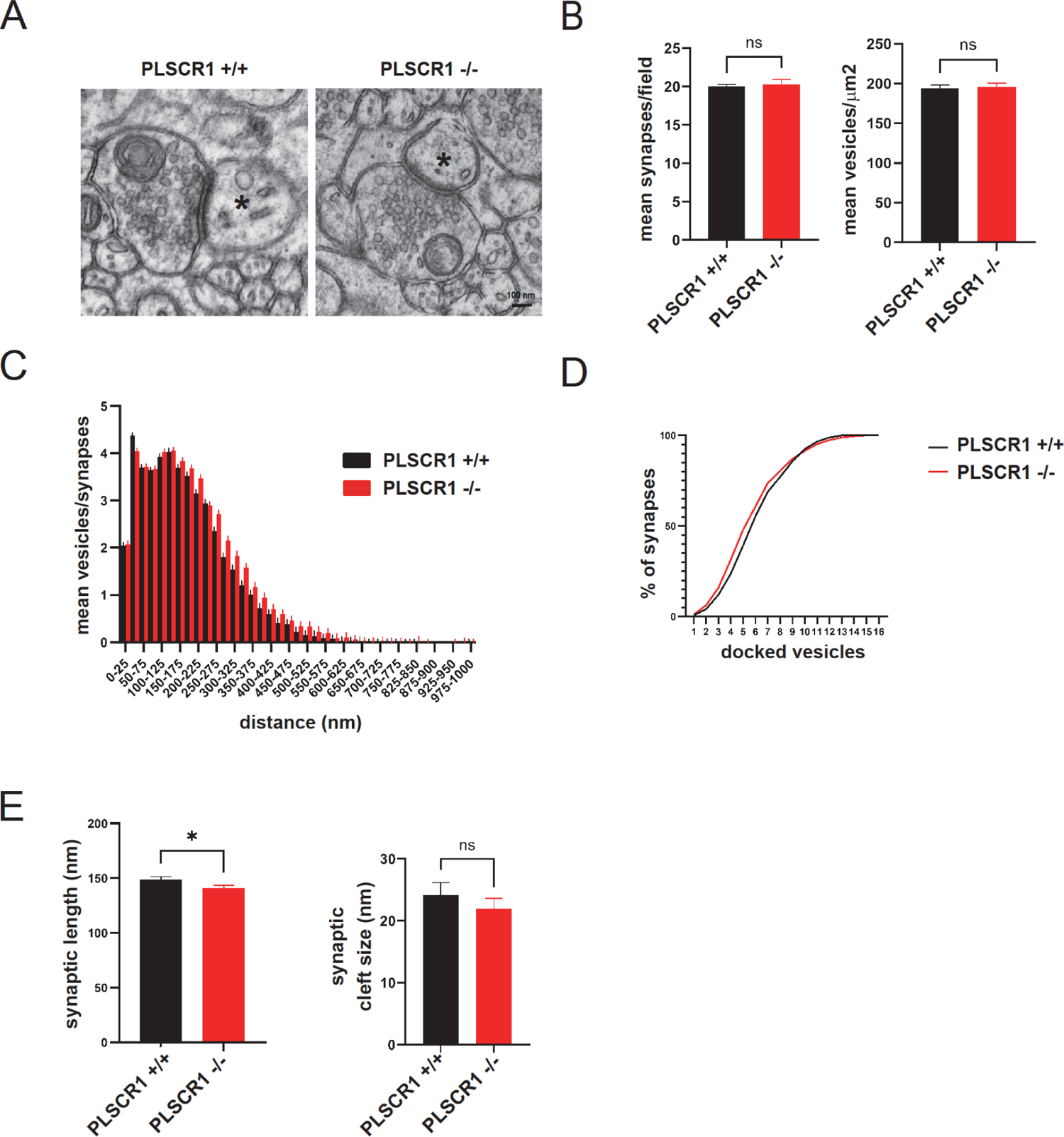
Morphometric analysis of cerebellum slices observed by transmission electron microscopy. **A**, representative electron micrographs of cerebellum from PLSCR1^+/+^ and PLSCR1^-/-^ mice showing synapses between GrC and Purkinje cell dendrites (PC, asterisks). Bar: 100 nm. **B**, synapses number in each field of view were counted (5 fields of 4 μm^2^, n=3 mice) and the number of synaptic vesicles (SVs) counted and reported on the surface of a bouton for PLSCR1^+/+^ and PLSCR1^-/-^ mice (± SEM). **C**, the shortest distance between SVs and the presynaptic membrane facing the postsynaptic density were calculated. The graph represents the distribution of mean number of SVs found according to their distance to the synapse. **D**, graph representing cumulative distribution of docked vesicles at GrC-PC synapse (distance <50 nm). **E**, the presynaptic and postsynaptic membranes were manually delineated. The length of the presynapse and the mean distance between the pre- and post-synapse were calculated (synaptic cleft size). Statistical significance was assessed using unpaired t-test or Kolmogorov-Smirnov test for distribution analysis. ns: non significant, * p<0.05. (p=0.79 and p=0.788 (B); p=0.7453 (C); p=0.996 (D); p=0.0162 and p=0.2157 (E)). More than 210 synapses were analyzed for each genotype.

### GrC stimulation triggers PLSCR1-dependent PS egress at synapses

Rapid activity-dependent transbilayer movement of PLs, mainly PS and PE, occurs in synaptosomes prepared from the electric organ of the electric ray, *Narke japonica* (Lee et al., 2000). We also demonstrated PLSCR1-dependent PS egress in chromaffin cells during exocytosis (Ory et al., 2013). We therefore asked whether stimulation of GrC could lead to disruption of the plasma membrane asymmetry, and whether PLSCR1 was required for that process. To detect PS exposed to the extracellular leaflet of the plasma membrane, living GrC were stimulated with a 50 mM KCl solution containing fluorescent AnnexinV (AnxA5), which selectively binds to PS (Andree et al., 1990). GrC were then fixed and stained for F-actin to delineate neuronal processes and to quantify AnxA5 staining. Compared to unstimulated neurons, depolarization of *Plscr1*^+/+^ GrC induced an increase in AnxA5 staining (Figure 3A, B). Interestingly, the increase in AnxA5 staining was abrogated in stimulated *Plscr1*^-/-^ GrC (Figure 3A,B) indicating that PLSCR1 was required for the activity-dependent translocation of PS to the extracellular leaflet.

**Figure 3:**
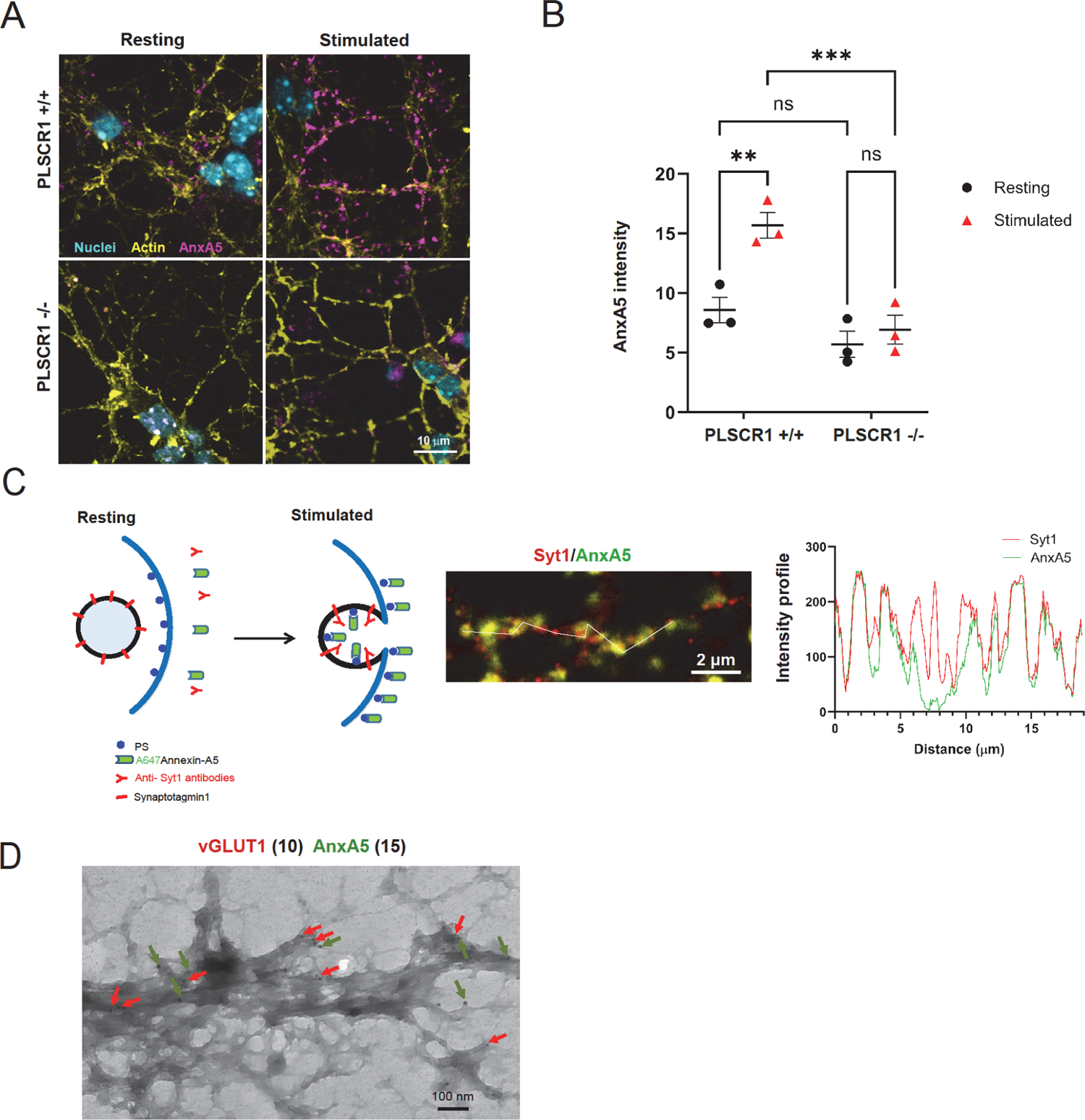
PLSCR1-dependent PS egress occurs at synapses. **A**, cultured GrC neurons from PLSCR1^+/+^ or PLSCR1^-/-^ mice were stimulated for 10 min with 59 mM K+ or maintained under resting condition in the presence of 1μg/ml fluorescent AnxA5 (magenta). Cells were fixed, counterstained for nuclei (blue) and actin (yellow) and observed under a confocal microscope. **B**, quantification of AnxA5 mean intensity of PLSCR1^+/+^ or PLSCR1^-/-^ neurons stimulated for 10 min with 59 mM K+ or maintained under resting condition (10 fields of view per experiments (±SEM), n=3 independent experiments). **C**, scheme of Syt1 staining assay. Living PLSCR1^+/+^ or PLSCR1^-/-^ GrC neurons were incubated in culture medium containing 30 mM KCl, anti-Syt1 antibodies directed against the luminal domain of Syt1 and fluorescent AnxA5 (green). Cells were fixed and anti-Syt1 antibody revealed with fluorescent secondary antibody without cell permeabilisation to reveal Syt1 at the plasma membrane (red). Cells were observed under a confocal microscope and intensity profile along the depicted line is shown. **D,** representative electron micrograph of plasma membrane sheets prepared from GrC neurons stimulated for 10 min in the presence of gold-conjugated AnxA5. Labeling of AnxA5 and vGlut1 were revealed with 15 nm (green arrows) and 10 nm (red arrows) gold particles respectively. Statistical significance was assessed using 2 way ANOVA with Holm-Sidak post hoc test. ns: non significant, **, p<0.01; ***, p<0.001. At least 21 fields of view from 3 independent experiments were analyzed for each condition.

AnxA5 staining organized as discrete spots along neuronal processes (Figure 3A) suggesting that PS egress could preferentially occur at the synapse. To test this hypothesis, we took advantage of anti-Syt1 antibodies that recognize the luminal domain of the integral SV protein Syt1, which is transiently accessible from the extracellular space upon SV fusion with the plasma membrane (Figure 3C). Incubating living GrC with both fluorescent AnxA5 and anti-Syt1 antibodies showed that AnxA5 and Syt1 stainings partially overlapped indicating that PS egress occurs at active synapses (Figure 3C). We also prepared plasma membrane sheets of stimulated neurons to determine more precisely where PS egress occurs. Thus, *Plscr1*^+/+^ GrC were stimulated in the presence of gold-coupled AnxA5 and the distribution of AnxA5 was compared to vGlut1 on membrane sheets. As shown in Figure 3D, clusters of AnxA5 gold particles were found in membrane patches close to vGlut1 staining. Altogether, these data indicate that activity-dependent PS egress occurs at synapses, and that this egress requires expression of PLSCR1.

### GFP-PLSCR1 restores PS egress in *Plscr1*^-/-^ GrCs

To confirm that PLSCR1 is essential for PS egress, we performed rescue experiments. GFP-tagged PLSCR1 protein was expressed in *Plscr1*^-/-^ GrC and PS egress in response to KCl-dependent stimulation was analyzed on plasma membrane sheets. We observed GFP-PLSCR1 in membrane patches containing both vGlut1 and gold-coupled AnxA5 in response to cell stimulation (Figure 4A). In addition, the percentage of membrane patches containing both vGlut1 and AnxA5 was higher in neurons expressing GFP-PLSCR1 than in non-transfected *Plscr1*^-/-^ neurons (Figure 4C). These data confirm that PLSCR1 is required for activity-dependent PS egress at synapses.

**Figure 4:**
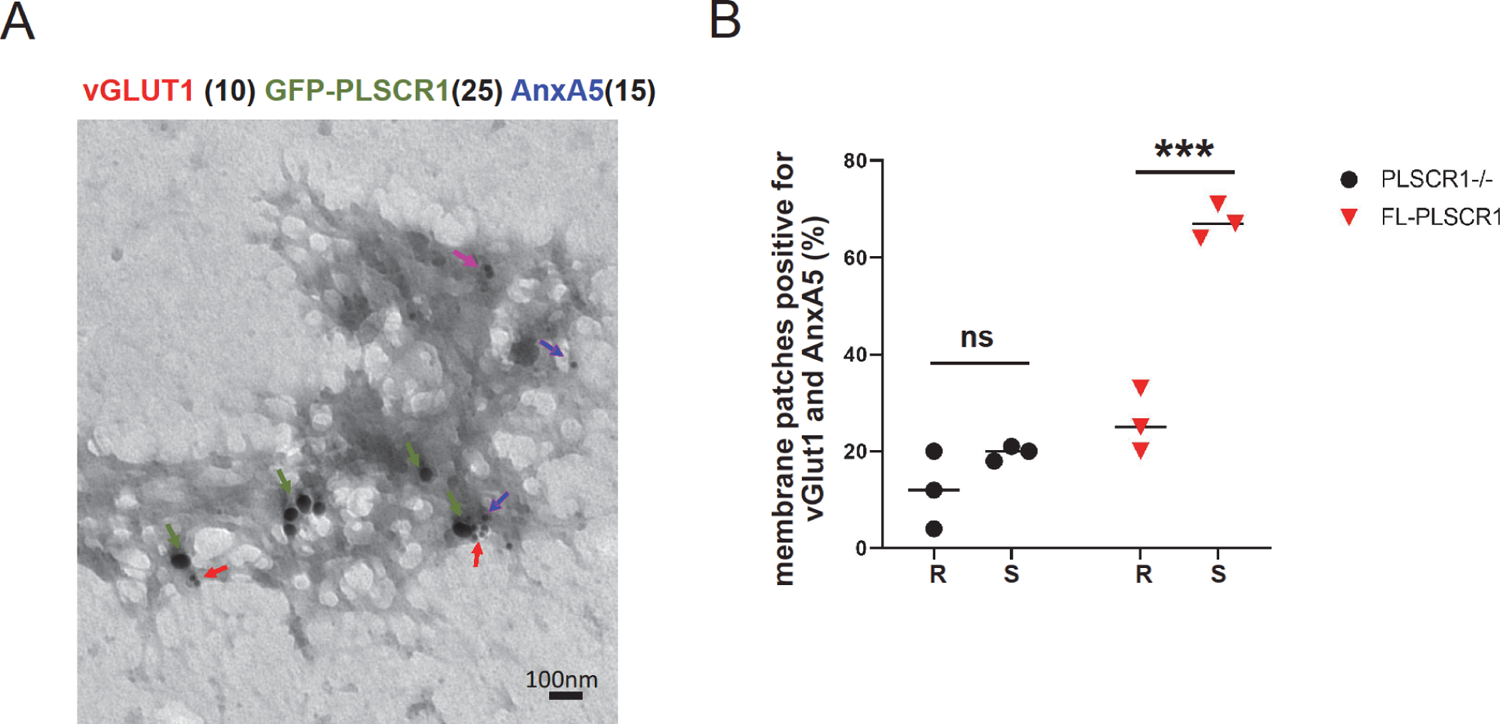
Overexpresion of PLSCR1 restores PS egress in PLSCR1^-/-^ GrC neurons. **A**, representative electron micrograph of plasma membrane sheets prepared from PLSCR1^-/-^ GrC neurons expressing GFP-PLSCR1. Neurons were stimulated for 10 min with 59 mM K^+^ (S) or maintained under resting condition (R) in the presence of gold-conjugated AnxA5 (15 nm beads). Immunolabeling of vGlut1 and GFP were revealed with 10 nm (red arrows), 15 nm (blue arrows) and 25 nm (green arrows) gold particles respectively. **B**, graph represents the % of synapses (membrane patches vGlut1 positive) closely associated to AnxA5 beads. Each point represent the mean of 3 independent experiments. In each experiment, 20 to 30 images per conditions were quantified. ns, non significant; ***, p<0.001

### PLSCR1 is required for evoked synaptic transmission

A key unsolved question is whether the PS egress controlled by PLSCR1 is involved in neurotransmission. In the cerebellar cortex, GrCs convey high-frequency information (several hundreds of Hz) to Purkinje cells (PCs) and molecular layer interneurons. GrC synapses stand out from most other synapse types by the striking ability to recruit reluctant vesicles within milliseconds to sustain the release of glutamate at such extreme frequencies. This phenomenon underlies a large facilitation of glutamate release during paired-pulse stimulation elicited at high frequency (Doussau et al., 2017; Miki et al., 2016). GrCs can also be endowed with specific mechanisms optimizing the recycling of used SVs as the high facilitation of glutamate release observed during high frequency trains is maintained for hundreds of milliseconds (Doussau et al., 2017). Hence, to test whether PLSCR1 is required for these presynaptic properties of GrCs, we prepared acute cerebellar slices from *Plscr1*^+/+^ and *Plscr1* ^-/-^ littermates in which we recorded excitatory postsynaptic currents (EPSCs) in PCs evoked by GrC axon stimulation (parallel fibres, PFs, Figure 5A) with either twin stimuli (paired-pulse stimulation at 50 Hz) or high frequency trains. As expected in *Plscr1*^+/+^ mice, amplitudes of EPSCs increased after the first stimulus (Figure 5B). Furthermore, excitatory neurotransmission was facilitated and this facilitation was maintained during tens of stimuli (Figure 5C). In contrast, in *Plscr1*^-/-^ mice, paired-pulse facilitation was strongly reduced (mean paired-pulse ratio: 2.03 ± 0.7 for *Plscr1*^+/+^ versus 1.59 ± 0.07 for *Plscr1*^-/-^, *t*-test, p = 0.001, n= 7 for *Plscr1*^+/+^ and n=8 for *Plscr1* ^-/-^). In addition, facilitation during the high frequency train was only transient, disappearing rapidly after the first ten stimuli (Figure 5C).

**Figure 5:**
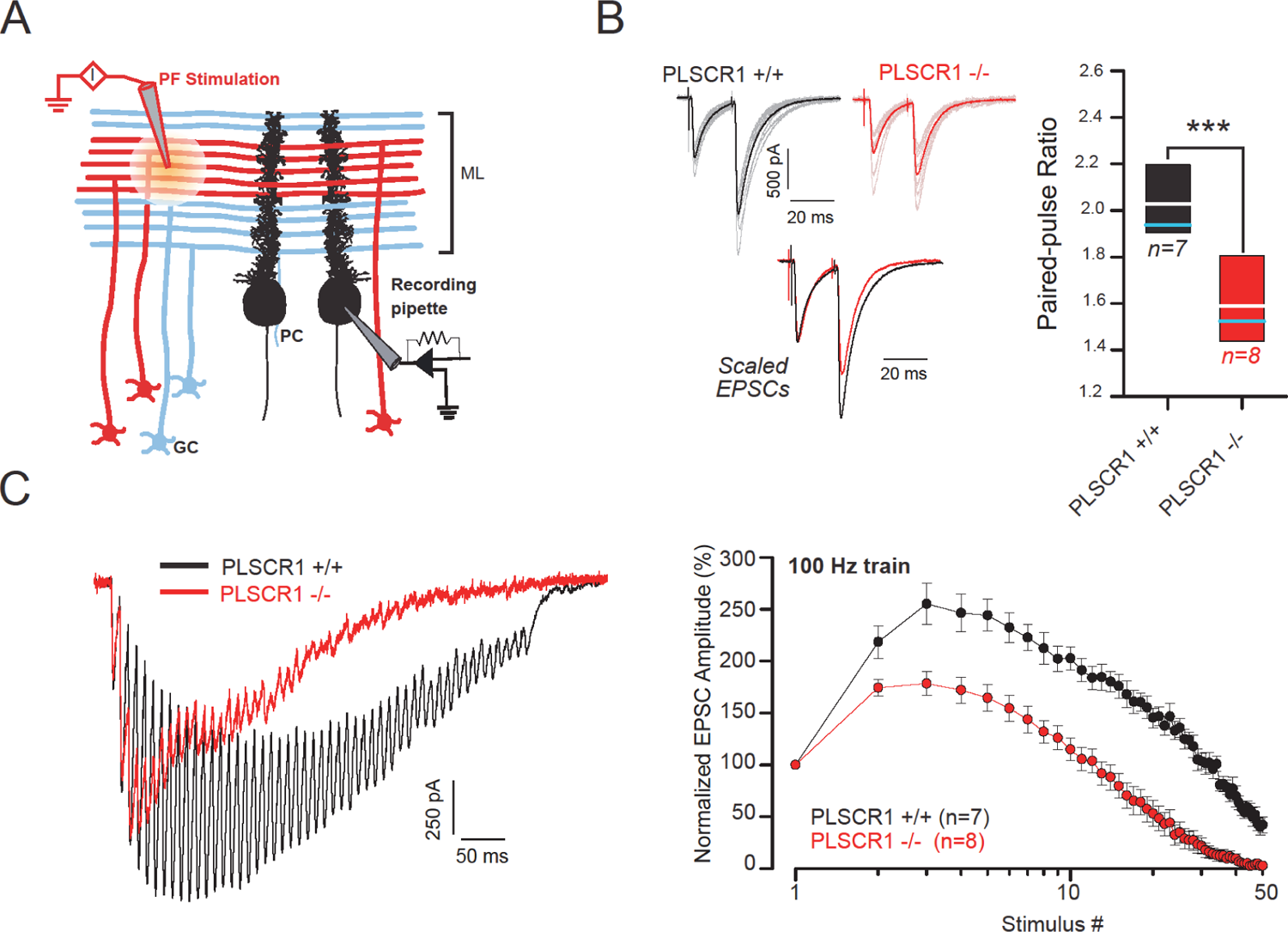
PLSCR1 is required to sustain synaptic transmission at high firing rate. **A**, schematic representing the connectivity in the cerebellar cortex and the position of the stimulating and recording electrodes. (GC: granule cell; PC: Purkinje cell, PF: parallel fiber, ML: molecular layer). **B,** *Left:* representative EPSCs evoked by paired-pulse stimulation (50 Hz) of PFs recorded in *Plscr1*^+/+^ and *Plscr1*^-/-^ mice acute slices (black and red traces respectively). Thick lines correspond to averaged EPSCs of at least 10 successive EPSCs (thin lines). Lower traces correspond to normalized averaged EPSCs. *Right*: Box-plots showing the values of the paired-pulse ratio obtained in *Plscr1*^+/+^ and *Plscr1*^-/-^ acute slice. White and blue lines correspond to mean and median values respectively (± SEM). **C,** *Left:* representative recording traces evoked by PF stimulations at 100 Hz (50 pulses) and recorded in *Plscr1*^+/+^ and *Plscr1*^-/-^ mice acute slices (black and red traces respectively). *Right:* Mean values of normalized EPSC amplitude elicited by trains of stimulation at 100 Hz and recorded in *Plscr1*^+/+^ and *Plscr1*^-/-^ mice acute slices (black and red traces respectively, ± SEM)

The absence of observable abnormalities in the ultrastructure of *Plscr1*^-/-^ GrC synapses (as shown in Figure 2) suggests that the presynaptic dysfunctions observed are unlikely to be caused by fewer synaptic vesicles in GrC boutons or major anatomical defects. Instead, it is more likely that PLSCR1 plays a key role in facilitating the rapid recruitment of reluctant vesicles (exocytosis) and/or in the recycling of previously used SVs (endocytosis).

### PLSCR1 is required for SV retrieval

To ascertain whether the effects observed on neurotransmission in the absence of PLSCR1 were due to altered SV exocytosis or endocytosis, we performed real-time monitoring of the genetically-encoded synaptophysin-pHluorin (Syp-pH) reporter. Syp-pH is widely used to report SV recycling since it responds to local pH changes occurring during SV exocytosis and endocytosis (Granseth et al., 2006; Jäpel et al., 2020; Nicholson-Fish et al., 2015). In resting conditions, as pHluorin faces the SV lumen, the fluorescence of Syp-pH is quenched by the acidic pH of the SV. Upon exocytosis, Syp-pH is exposed to the neutral extracellular medium, resulting in fluorescence unquenching, the quantification of which represents an assessment of the extent of synaptic exocytosis. After termination of stimulation, Syp-pH fluorescence intensity decreases rapidly as it is retrieved by the endocytic SVs, which quickly reacidify. As SV endocytosis is the rate limiting step in this fluorescence decrease, measurement of post-stimulus Syp-pH fluorescence decay is a reliable indicator of compensatory endocytosis (Atluri and Ryan, 2006; Egashira et al., 2015; Granseth et al., 2006; Sankaranarayanan and Ryan, 2000). *Plscr1*^+/+^ or *Plscr1*^-/-^ GrCs were transfected with Syp-pH and stimulated with a train of high-frequency action potentials (40Hz, 10s) to evoke robust exocytosis (Nicholson-Fish et al., 2015). Analysis of the fluorescence intensity profiles showed a fast rise in fluorescence upon stimulation for both genotypes (Figure 6A, B) and a progressive return to the fluorescence baseline for *Plscr1*^+/+^ GrCs. Although no significant differences in the amount of Syp-pH visiting the cell surface was observed (Figure 6C), fluorescence decay was severely delayed in *Plscr1*^-/-^ GrCs (Figure 6B, D), suggesting that SV exocytosis was unaltered, but that endocytosis was impaired in the absence of PLSCR1. To confirm the requirement for PLSCR1 in SV endocytosis, we performed restoration experiments in *Plscr1*^-/-^ GrCs using exogenous expression of the mCherry-tagged version of PLSCR1. Expression of exogenous PLSCR1 did not affect the stimulation evoked rise in fluorescence but significantly restored the decay of fluorescence intensity (Figure 6B, D), in line with the idea that PLSCR1 is involved in compensatory endocytosis.

**Figure 6:**
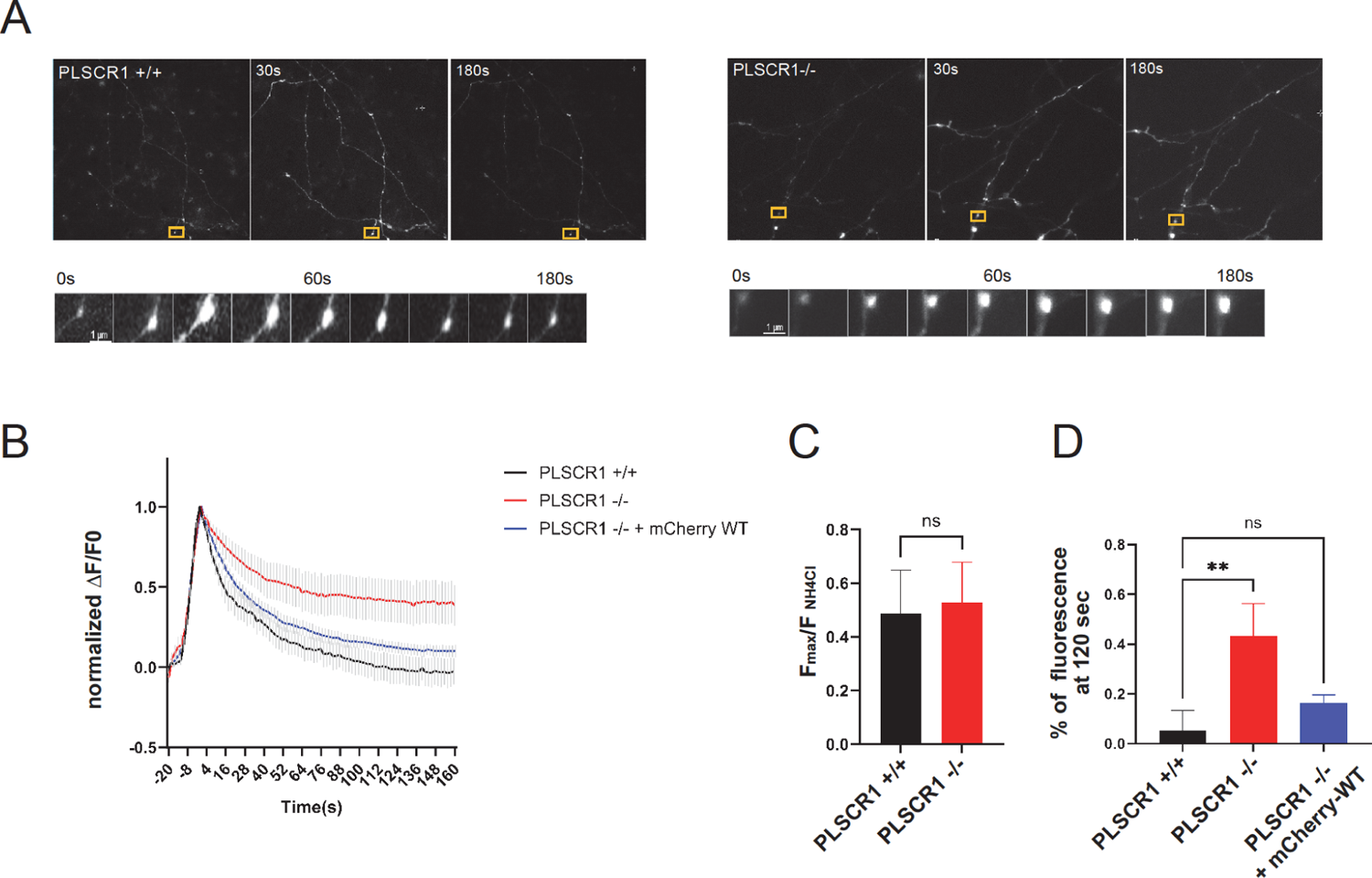
PLSCR1 is required for efficient SVs endocytosis during high frequency stimulation of GrC neurons. **A**, representative images of GrC neuron transfected with Synaptophysin-pHluorin (Syp-pH) and stimulated with a train of 400 action potentials delivered at 40 Hz. Still images show *Plscr1*^+/+^ and *Plscr1*^-/-^ GrC neurons before stimulation, 30 s and 180 s after stimulation. Inset shows a representative time course of Syp-pH variation at a single synapse. **B**, average normalized time course of fluorescence response of Syp-pH represented as ΔF/F0 +/- SEM (5 independent preparations, n≥15 neurons by condition) in GrC neurons *Plscr1*^+/+^, *Plscr1*^-/-^ and *Plscr1*^-/-^ transfected with mCherry-PLSCR1. **C**, mean peak of Syp-pH response during stimulation. Maximum Syp-pH intensity after stimulation was normalized to the total amount of Syp-pH at synapse obtained after incubation of neurons with 50 mM NH_4_Cl (± SEM) **D**, mean normalized intensity of Syp-pH fluorescence remaining after 2 min imaging (± SEM). ns: non significant; **, p<0.01

## Discussion

To sustain neurotransmission during elevated neuronal activity, a major reorganization of the neuronal plasma membrane has to occur to integrate the SV at the exocytic site and to subsequently retrieve lipids and proteins to replenish the vesicular pool (Binotti et al., 2021). While experimental evidence points to a major role for phosphoinositides and their metabolism in this process, the role of structural phospholipids such as PS and the regulation of their localisation has remained unexplored. Here, we provide the first evidence that PLSCR1 randomizes PS at synapses during heightened neuronal activity and that the resulting loss of plasma membrane asymmetry is dispensable for SVs exocytosis but necessary for compensatory SV endocytosis. As a consequence of impaired vesicular pool replenishment, synaptic transmission at GrC to PC synapses in cerebellar acute slices displays accelerated depletion in the absence of PLSCR1. Therefore, we propose that PLSCR1-dependent PS egress controls compensatory endocytosis in GrC and contributes to sustain neurotransmission at high frequencies.

### PLSCR1 and plasma membrane asymmetry disruption

Immunofluorescence and immunogold electron microscopy approaches revealed that, 1) PLSCR1 is enriched at synaptic terminals, and 2) PLSCR1 is required for activity-dependent PS egress at these synapses. This is the first evidence that PLSCR1 promotes shuffling of the structural lipid PS in neurons. PLSCR1 has been involved in structural lipid mixing both *in vitro* (Rayala et al., 2014) and in immune and chromaffin cells during regulated exocytosis (Kato et al., 2002; Ory et al., 2013). Our observation that PS egress is restricted to synapses indicates that PLSCR1 is probably activated locally as has been reported in chromaffin cells (Ory et al., 2013). In these cells, PS egress occurred only in the vicinity of exocytosis sites, despite PLSCR1 being homogeneously distributed at the plasma membrane (Ory et al., 2013). The low affinity of PLSCR1 for Ca^2+^ (Stout et al., 1998) together with the activity described here occurring only after neurotransmission suggests that PLSCR1 may only be activated when intracellular Ca^2+^ reaches a critical concentration. Typically, low affinity Ca^2+^ sensors must be localised close to the exocytic site, since the Ca^2+^ burst is spatially constrained within a microdomain at the presynaptic active zone (Neher and Sakaba, 2008). Our finding that PLSCR1 is required for SV endocytosis during periods of high activity, suggests that its activity-dependent triggering of PS redistribution must occur in or directly adjacent to the active zone. This latter region is termed the periactive zone, and is where SV endocytosis is proposed to occur (Gad et al., 1998; Watanabe et al., 2013). It is particularly relevant in high frequency synapses, where rapid and repetitive stimulation leads to the accumulation of intracellular Ca^2+^ (Delvendahl et al., 2015). Thus, the localized activation of PLSCR1 and the ensuing PS redistribution play a crucial role in modulating synaptic function, especially in scenarios involving high-frequency synaptic activity.

### How can the local loss of plasma membrane asymmetry control compensatory endocytosis?

In the absence of PLSCR1, PS translocation to the extracellular leaflet is blocked and SV endocytosis, as measured by the retrieval of Syp-pH, is retarded dramatically. What could the requirement for PS egress be in SV endocytosis? The inner leaflet of the plasma membrane is negatively charged, due to the presence of anionic phospholipids including phosphatidic acid (PA), phosphoinositides (PIs) and PS. Compared to PA and PIs, which represent only a minor fraction of the phospholipids present at the plasma membrane, PS is the predominant phospholipid, and its redistribution from one leaflet to the other should alter the biophysical properties of the plasma membrane (Puchkov and Haucke, 2013; Yeung et al., 2008). Depending on the timing of lipid scrambling relative to SV fusion, a local decrease of negative charges by local PS depletion may, for example, reduce the repulsive forces of opposing membrane to favor fusion between SVs and the plasma membrane (Davletov and Montecucco, 2010). However, the amount of Syp-pH visiting the plasma membrane during activity was unaffected in *Plscr1*^-/-^ GrCs, suggesting that exocytosis was unaltered. In agreement, large dense core vesicle fusion was unaltered in chromaffin cells from *Plscr1*^-/-^ mice (Ory et al., 2013).

An alteration in local membrane charge is not predicted to impact SV endocytosis. This is because accumulation of negatively charged phospholipids such as PtdIns(4,5)P_2_ increase the recruitment of cytosolic proteins with positively charged residues (Clarke et al., 2020; Kay and Fairn, 2019) to favor membrane bending (Micheva et al., 2001; Puchkov and Haucke, 2013). In contrast, PS egress may modify plasma membrane fluidity. Enriched in tightly packed sphingolipids, the outer plasma membrane leaflet is highly ordered and rigid (Gupta et al., 2020; Lorent et al., 2020). Insertion of SVs that are highly enriched in cholesterol will most likely modify the lipid organization of the presynaptic plasma membrane (Takamori et al., 2006). Indeed, membrane cholesterol content affects lipid lateral self-diffusion, confines SV proteins and limits their diffusive behavior (Byczkowicz et al., 2018; Dason et al., 2014; Mercer et al., 2011; Wilson et al., 2020). PS and cholesterol are co-distributed in the inner leaflet of the plasma membrane and the local decrease of PS may also lead to concomitant decrease in cholesterol content (Maekawa and Fairn, 2015) helping to fluidize the leaflet and clear the active zone. Alternatively, changes in PS distribution may unlock protein function. For example, PS shapes the transmembrane domain of synaptogyrin to bend membrane and generate SV of homogeneous size. To achieve this, PS must be present simultaneously on both leaflets to induce a structural change in synaptogyrin, a function that is intrinsic to PLSCR1 (Yu et al., 2023). It remains to be seen whether proteins involved in endocytosis may have the same properties and the precise role of phospholipids egress needs to be further explored.

### PLSCR1 and GrC neurotransmission

Synaptic plasticity allows for fast adaptation of synaptic strength to neuronal network activity, which is critical for information processing. This comes in two main modes: 1) functional plasticity, which changes the strength of the synapse by modifying signal transmission and 2) structural plasticity, which alters the number and/or shape of synapses, resulting in differential connectivity between neurons (Caroni et al., 2012). Morphometric analysis of *Plscr1*^-/-^ cerebellum slices showed no major alteration within presynaptic terminals, suggesting that PLSCR1 may impair functional, rather than structural, plasticity. In the cerebellar cortex, GrC-PCs synapses have to transmit sensorimotor information elicited at extreme frequencies (several hundred of Hz to kHz, Rancz et al., 2007). To do so, these synapses are endowed with specific presynaptic processes allowing a fast and sustained boost (facilitation) of the release neurotransmitters during bursts of high-frequency activities. The strikingly large facilitation that occurs during paired-pulse facilitation and during the first phase of high-frequency trains is underpinned by the ultra-fast recruitment of a pool of reluctant SVs that can only be mobilized by high-frequency activities (Doussau et al., 2017; Miki et al., 2016). This phenomenon is affected in *Plscr1*^-/-^ GrC terminals, indicating that PLSCR1 activity control directly or indirectly the size of the pool of reluctant pool and/or its recruitment. The presence of PLSCR1 is also required for the fast mobilization of recycling SVs that support the maintenance of a high level of facilitation during prolonged stimulations. Whether SV mobilization defects are directly or indirectly related to the slowing down of SV endocytosis need to be clarified. Furthermore, we cannot exclude alternative mechanisms underlying the facilitation defect as altered clearance of SV cargo from release sites or an increase in release probability resulting from compensatory mechanisms.

Interestingly, we detected PLSCR1 in the cerebellum and the olfactory bulb, two brain regions known to sustain high frequency stimulation. In contrast, PLSCR1 is barely detected in other brain area. PLSCR1 appears therefore a good candidate to play a specific role in high-frequency synaptic transmission by scrambling PLs at synapse, a fast way to modify plasma membrane properties.

## Acknowledgements

This work was financially supported by the Agence Nationale pour la Recherche (“LipidTrans4NeuroTraffic”, N° ANR-19-CE16-0012-01) to SG; by a fellowship from la Fondation pour la Recherche Médicale (n° FDT202106013135) and a travel grant from The Company of Biologists (n° JCSTF1911344) to MC; and by the Wellcome Trust Investigator Award to MAC (204954/Z/16/Z). For open access, authors have applied a CC-BY public copyright license to any accepted manuscript version arising from this submission. INSERM is providing salary to MFB, SCG, SG and NV. We acknowledge the Plateforme Imagerie In Vitro at CNRS UPS3256, Dr Jean-Daniel Fauny (IBMC, UPR3512, Strasbourg) for technical assistance with live cell imaging microscopy set up and the animal facility of Institut des Neurosciences Cellulaires et Intégratives (Chronobiotron UMS 3415). The authors are grateful to Charlotte Caquineau and Audrey Groh for technical assistance.

## References

Amir-Moazami, O., Alexia, C., Charles, N., Launay, P., Monteiro, R. C. and Benhamou, M. (2008). Phospholipid Scramblase 1 Modulates a Selected Set of IgE Receptor-mediated Mast Cell Responses through LAT-dependent Pathway. Journal of Biological Chemistry 283, 25514– 25523.

Andree, H. A., Reutelingsperger, C. P., Hauptmann, R., Hemker, H. C., Hermens, W. T. and Willems, G. M. (1990). Binding of vascular anticoagulant alpha (VAC alpha) to planar phospholipid bilayers. Journal of Biological Chemistry 265, 4923–4928.

Atluri, P. P. and Ryan, T. A. (2006). The Kinetics of Synaptic Vesicle Reacidification at Hippocampal Nerve Terminals. J. Neurosci. 26, 2313–2320.

Audo, R., Hua, C., Hahne, M., Combe, B., Morel, J. and Daien, C. I. (2017). Phosphatidylserine Outer Layer Translocation Is Implicated in IL-10 Secretion by Human Regulatory B Cells. PLOS ONE 12, e0169755.

Bassé, F., Stout, J. G., Sims, P. J. and Wiedmer, T. (1996). Isolation of an Erythrocyte Membrane Protein that Mediates Ca2+-dependent Transbilayer Movement of Phospholipid. J. Biol. Chem. 271, 17205–17210.

Bevers, E. M. and Williamson, P. L. (2016). Getting to the Outer Leaflet: Physiology of Phosphatidylserine Exposure at the Plasma Membrane. Physiological Reviews 96, 605–645.

Binotti, B., Jahn, R. and Pérez-Lara, Á. (2021). An overview of the synaptic vesicle lipid composition. Archives of Biochemistry and Biophysics 709, 108966.

Bolz, S., Kaempf, N., Puchkov, D., Krauss, M., Russo, G., Soykan, T., Schmied, C., Lehmann, M., Müller, R., Schultz, C., et al. (2023). Synaptotagmin 1-triggered lipid signaling facilitates coupling of exo- and endocytosis. Neuron 0,.

Bonnycastle, K., Davenport, E. C. and Cousin, M. A. (2020). Presynaptic dysfunction in neurodevelopmental disorders: Insights from the synaptic vesicle life cycle. Journal of Neurochemistry n/a,.

Byczkowicz, N., Ritzau-Jost, A., Delvendahl, I. and Hallermann, S. (2018). How to maintain active zone integrity during high-frequency transmission. Neuroscience Research 127, 61–69.

Caroni, P., Donato, F. and Muller, D. (2012). Structural plasticity upon learning: regulation and functions. Nat Rev Neurosci 13, 478–490.

Chanaday, N. L., Cousin, M. A., Milosevic, I., Watanabe, S. and Morgan, J. R. (2019). The Synaptic Vesicle Cycle Revisited: New Insights into the Modes and Mechanisms. J. Neurosci. 39, 8209– 8216.

Cheung, G. and Cousin, M. A. (2011). Quantitative analysis of synaptic vesicle pool replenishment in cultured cerebellar granule neurons using FM dyes. J Vis Exp 3143.

Clarke, R. J., Hossain, K. R. and Cao, K. (2020). Physiological roles of transverse lipid asymmetry of animal membranes. Biochimica et Biophysica Acta (BBA) - Biomembranes 1862, 183382.

Daleke, D. L. (2003). Regulation of transbilayer plasma membrane phospholipid asymmetry. J. Lipid Res. 44, 233–242.

Dason, J. S., Smith, A. J., Marin, L. and Charlton, M. P. (2014). Cholesterol and F-actin are required for clustering of recycling synaptic vesicle proteins in the presynaptic plasma membrane. The Journal of Physiology 592, 621–633.

Davletov, B. and Montecucco, C. (2010). Lipid function at synapses. Current Opinion in Neurobiology 20, 543–549.

Delavoie, F., Royer, C., Gasman, S., Vitale, N. and Chasserot-Golaz, S. (2021). Transmission Electron MicroscopyMicroscopy and TomographyTomography on Plasma MembranePlasma membranes Sheets to Study Secretory Docking. In Exocytosis and Endocytosis: Methods and Protocols (ed. Niedergang, F.), Vitale, N.), and Gasman, S.), pp. 301–309. New York, NY: Springer US.

Delvendahl, I., Jablonski, L., Baade, C., Matveev, V., Neher, E. and Hallermann, S. (2015). Reduced endogenous Ca^2+^ buffering speeds active zone Ca^2+^ signaling. Proceedings of the National Academy of Sciences 112, E3075–E3084.

D’Mello, S. R., Galli, C., Ciotti, T. and Calissano, P. (1993). Induction of apoptosis in cerebellar granule neurons by low potassium: inhibition of death by insulin-like growth factor I and cAMP. PNAS 90, 10989–10993.

Doussau, F., Schmidt, H., Dorgans, K., Valera, A. M., Poulain, B. and Isope, P. (2017). Frequency-dependent mobilization of heterogeneous pools of synaptic vesicles shapes presynaptic plasticity. eLife 6, e28935.

Dunkley, P. R., Jarvie, P. E. and Robinson, P. J. (2008a). A rapid Percoll gradient procedure for preparation of synaptosomes. Nat Protoc 3, 1718–1728.

Dunkley, P. R., Jarvie, P. E. and Robinson, P. J. (2008b). A rapid Percoll gradient procedure for preparation of synaptosomes. Nat Protoc 3, 1718–1728.

Egashira, Y., Takase, M. and Takamori, S. (2015). Monitoring of Vacuolar-Type H+ ATPase-Mediated Proton Influx into Synaptic Vesicles. J. Neurosci. 35, 3701–3710.

Gabel, M., Delavoie, F., Demais, V., Royer, C., Bailly, Y., Vitale, N., Bader, M.-F. and Chasserot-Golaz, S. (2015). Annexin A2–dependent actin bundling promotes secretory granule docking to the plasma membrane and exocytosis. The Journal of Cell Biology 210, 785–800.

Gad, H., Löw, P., Zotova, E., Brodin, L. and Shupliakov, O. (1998). Dissociation between Ca2+-triggered synaptic vesicle exocytosis and clathrin-mediated endocytosis at a central synapse. Neuron 21, 607–616.

Granseth, B., Odermatt, B., Royle, S. J. and Lagnado, L. (2006). Clathrin-Mediated Endocytosis Is the Dominant Mechanism of Vesicle Retrieval at Hippocampal Synapses. Neuron 51, 773–786.

Gupta, A., Korte, T., Herrmann, A. and Wohland, T. (2020). Plasma membrane asymmetry of lipid organization: fluorescence lifetime microscopy and correlation spectroscopy analysis[S]. Journal of Lipid Research 61, 252–266.

Jäpel, M., Gerth, F., Sakaba, T., Bacetic, J., Yao, L., Koo, S.-J., Maritzen, T., Freund, C. and Haucke, V. (2020). Intersectin-Mediated Clearance of SNARE Complexes Is Required for Fast Neurotransmission. Cell Reports 30, 409–420.e6.

Kato, N., Nakanishi, M. and Hirashima, N. (2002). Transbilayer Asymmetry of Phospholipids in the Plasma Membrane Regulates Exocytotic Release in Mast Cells. Biochemistry 41, 8068–8074.

Kay, J. G. and Fairn, G. D. (2019). Distribution, dynamics and functional roles of phosphatidylserine within the cell. Cell Commun Signal 17, 1–8.

Kobayashi, T. and Menon, A. K. (2018). Transbilayer lipid asymmetry. Current Biology 28, R386– R391.

Krämer, D. and Minichiello, L. (2010). Cell Culture of Primary Cerebellar Granule Cells. In Mouse Cell Culture (ed. Ward, A.) and Tosh, D.), pp. 233–239. Totowa, NJ: Humana Press.

Lawrie, A. M., Graham, M. E., Thorn, P., Gallacher, D. V. and Burgoyne, R. D. (1993). Synchronous calcium oscillations in cerebellar granule cells in culture mediated by NMDA receptors: NeuroReport 4, 539–542.

Lee, D.-S., Hirashima, N. and Kirino, Y. (2000). Rapid transbilayer phospholipid redistribution associated with exocytotic release of neurotransmitters from cholinergic nerve terminals isolated from electric ray Narke japonica. Neuroscience Letters 291, 21–24.

Lein, E. S., Hawrylycz, M. J., Ao, N., Ayres, M., Bensinger, A., Bernard, A., Boe, A. F., Boguski, M. S., Brockway, K. S., Byrnes, E. J., et al. (2007). Genome-wide atlas of gene expression in the adult mouse brain. Nature 445, 168–176.

Lorent, J. H., Levental, K. R., Ganesan, L., Rivera-Longsworth, G., Sezgin, E., Doktorova, M., Lyman, E. and Levental, I. (2020). Plasma membranes are asymmetric in lipid unsaturation, packing and protein shape. Nature Chemical Biology 16, 644–652.

Maekawa, M. and Fairn, G. D. (2015). Complementary probes reveal that phosphatidylserine is required for the proper transbilayer distribution of cholesterol. J Cell Sci 128, 1422–1433.

Malacombe, M., Ceridono, M., Calco, V., Chasserot-Golaz, S., McPherson, P. S., Bader, M.-F. and Gasman, S. (2006). Intersectin-1L nucleotide exchange factor regulates secretory granule exocytosis by activating Cdc42. The EMBO Journal 25, 3494–3503.

Manich, M., Boquet-Pujadas, A., Dallongeville, S., Guillen, N. and Olivo-Marin, J.-C. (2020). A Protocol to Quantify Cellular Morphodynamics: From Cell Labelling to Automatic Image Analysis. In Eukaryome Impact on Human Intestine Homeostasis and Mucosal Immunology (ed. Guillen, N.), pp. 351–367. Cham: Springer International Publishing.

Maritzen, T. and Haucke, V. (2018). Coupling of exocytosis and endocytosis at the presynaptic active zone. Neuroscience Research 127, 45–52.

Martin, S., Pombo, I., Poncet, P., David, B., Arock, M. and Blank, U. (2000). Immunologic Stimulation of Mast Cells Leads to the Reversible Exposure of Phosphatidylserine in the Absence of Apoptosis. Int Arch Allergy Immunol 123, 249–258.

Mercer, A. J., Chen, M. and Thoreson, W. B. (2011). Lateral mobility of presynaptic L-Type calcium channels at photoreceptor ribbon synapses. Journal of Neuroscience 31, 4397–4406.

Micheva, K. D., Holz, R. W. and Smith, S. J. (2001). Regulation of presynaptic phosphatidylinositol 4,5-biphosphate by neuronal activity. Journal of Cell Biology 154, 355–368.

Miki, T., Malagon, G., Pulido, C., Llano, I., Neher, E. and Marty, A. (2016). Actin- and Myosin-Dependent Vesicle Loading of Presynaptic Docking Sites Prior to Exocytosis. Neuron 91, 808– 823.

Neher, E. and Sakaba, T. (2008). Multiple Roles of Calcium Ions in the Regulation of Neurotransmitter Release. Neuron 59, 861–872.

Nicholson-Fish, J. C., Kokotos, A. C., Gillingwater, T. H., Smillie, K. J. and Cousin, M. A. (2015). VAMP4 Is an Essential Cargo Molecule for Activity-Dependent Bulk Endocytosis. Neuron 88, 973–984.

Ory, S., Ceridono, M., Momboisse, F., Houy, S., Chasserot-Golaz, S., Heintz, D., Calco, V., Haeberle, A.-M., Espinoza, F. A., Sims, P. J., et al. (2013). Phospholipid Scramblase-1-Induced Lipid Reorganization Regulates Compensatory Endocytosis in Neuroendocrine Cells. J. Neurosci. 33, 3545–3556.

Puchkov, D. and Haucke, V. (2013). Greasing the synaptic vesicle cycle by membrane lipids. Trends in Cell Biology 23, 493–503.

Rancz, E. A., Ishikawa, T., Duguid, I., Chadderton, P., Mahon, S. and Häusser, M. (2007). High-fidelity transmission of sensory information by single cerebellar mossy fibre boutons. Nature 450, 1245–1248.

Rayala, S., Francis, V. G., Sivagnanam, U. and Gummadi, S. N. (2014). N-terminal Proline-rich Domain Is Required for Scrambling Activity of Human Phospholipid Scramblases. J. Biol. Chem. 289, 13206–13218.

Rysavy, N. M., Shimoda, L. M. N., Dixon, A. M., Speck, M., Stokes, A. J., Turner, H. and Umemoto, E. Y. (2014). Beyond apoptosis: The mechanism and function of phosphatidylserine asymmetry in the membrane of activating mast cells. BioArchitecture 4, 127–137.

Sankaranarayanan, S. and Ryan, T. A. (2000). Real-time measurements of vesicle-SNARE recycling in synapses of the central nervous system. Nat Cell Biol 2, 197–204.

Sjöstedt, E., Zhong, W., Fagerberg, L., Karlsson, M., Mitsios, N., Adori, C., Oksvold, P., Edfors, F., Limiszewska, A., Hikmet, F., et al. (2020). An atlas of the protein-coding genes in the human, pig, and mouse brain. Science 367, eaay5947.

Smrž, D., Lebduška, P., Dráberová, L., Korb, J. and Dráber, P. (2008). Engagement of Phospholipid Scramblase 1 in Activated Cells: IMPLICATION FOR PHOSPHATIDYLSERINE EXTERNALIZATION AND EXOCYTOSIS. Journal of Biological Chemistry 283, 10904–10918.

Stout, J. G., Zhou, Q., Wiedmer, T. and Sims, P. J. (1998). Change in Conformation of Plasma Membrane Phospholipid Scramblase Induced by Occupancy of Its Ca ^2+^ Binding Site. Biochemistry 37, 14860–14866.

Suzuki, J., Umeda, M., Sims, P. J. and Nagata, S. (2010). Calcium-dependent phospholipid scrambling by TMEM16F. Nature 468, 834–838.

Suzuki, J., Denning, D. P., Imanishi, E., Horvitz, H. R. and Nagata, S. (2013). Xk-Related Protein 8 and CED-8 Promote Phosphatidylserine Exposure in Apoptotic Cells. Science 341, 403–406.

Takamori, S., Holt, M., Stenius, K., Lemke, E. A., Grønborg, M., Riedel, D., Urlaub, H., Schenck, S., Brügger, B., Ringler, P., et al. (2006). Molecular Anatomy of a Trafficking Organelle. Cell 127, 831–846.

Vitale, N., Caumont, A.-S., Chasserot-Golaz, S., Du, G., Wu, S., Sciorra, V. A., Morris, A. J., Frohman, M. A. and Bader, M.-F. (2001). Phospholipase D1: a key factor for the exocytotic machinery in neuroendocrine cells. The EMBO Journal 20, 2424–2434.

Watanabe, S., Rost, B. R., Camacho-Pérez, M., Davis, M. W., Söhl-Kielczynski, B., Rosenmund, C. and Jorgensen, E. M. (2013). Ultrafast endocytosis at mouse hippocampal synapses. Nature 504, 242–247.

Wiedmer, T., Zhou, Q., Kwoh, D. Y. and Sims, P. J. (2000). Identification of three new members of the phospholipid scramblase gene family. Biochimica et Biophysica Acta (BBA) - Biomembranes 1467, 244–253.

Wilson, B. S., Pfeiffer, J. R. and Oliver, J. M. (2000). Observing Fcεri Signaling from the Inside of the Mast Cell Membrane. Journal of Cell Biology 149, 1131–1142.

Wilson, K. A., MacDermott-Opeskin, H. I., Riley, E., Lin, Y. and O’Mara, M. L. (2020). Understanding the Link between Lipid Diversity and the Biophysical Properties of the Neuronal Plasma Membrane. Biochemistry 59, 3010–3018.

Yang, H., Kim, A., David, T., Palmer, D., Jin, T., Tien, J., Huang, F., Cheng, T., Coughlin, S. R., Jan, Y. N., et al. (2012). TMEM16F Forms a Ca2+-Activated Cation Channel Required for Lipid Scrambling in Platelets during Blood Coagulation. Cell 151, 111–122.

Yeung, T., Gilbert, G. E., Shi, J., Silvius, J., Kapus, A. and Grinstein, S. (2008). Membrane Phosphatidylserine Regulates Surface Charge and Protein Localization. Science 319, 210–213.

Yu, T., Flores-Solis, D., Eastep, G. N., Becker, S. and Zweckstetter, M. (2023). Phosphatidylserine-dependent structure of synaptogyrin remodels the synaptic vesicle membrane. Nat Struct Mol Biol 30, 926–934.

Zachowski, A. (1993). Phospholipids in animal eukaryotic membranes: transverse asymmetry and movement. Biochemical Journal 294, 1–14.

Zhao, S., Ting, J. T., Atallah, H. E., Qiu, L., Tan, J., Gloss, B., Augustine, G. J., Deisseroth, K., Luo, M., Graybiel, A. M., et al. (2011). Cell type–specific channelrhodopsin-2 transgenic mice for optogenetic dissection of neural circuitry function. Nature Methods 8, 745–752.

Zhou, Q., Zhao, J., Stout, J. G., Luhm, R. A., Wiedmer, T. and Sims, P. J. (1997). Molecular Cloning of Human Plasma Membrane Phospholipid Scramblase A PROTEIN MEDIATING TRANSBILAYER MOVEMENT OF PLASMA MEMBRANE PHOSPHOLIPIDS. J. Biol. Chem. 272, 18240–18244.

Zhou, Q., Zhao, J., Wiedmer, T. and Sims, P. J. (2002). Normal hemostasis but defective hematopoietic response to growth factors in mice deficient in phospholipid scramblase 1. Blood 99, 4030–4038.

Zhou, Q., Ben-Efraim, I., Bigcas, J.-L., Junqueira, D., Wiedmer, T. and Sims, P. J. (2005). Phospholipid Scramblase 1 Binds to the Promoter Region of the Inositol 1,4,5-Triphosphate Receptor Type 1 Gene to Enhance Its Expression. Journal of Biological Chemistry 280, 35062–35068.

